# Museomics reveals the phylogenetic position of an enigmatic vertebrate family (Lophiiformes: Lophichthyidae)

**DOI:** 10.1101/2025.02.25.640136

**Authors:** Moritz Muschick, Lukas Rüber, Michael Matschiner

## Abstract

*Lophichthys boschmai*, the only member of the family Lophichthyidae, is an elusive lophiiform fish whose phylogenetic associations are debated. This family is among the last three vertebrate families lacking DNA sequence data, primarily due to the species’ rarity, lack of fresh material and challenges associated with sequencing formalin-treated museum specimens. In this study, we provide mitochondrial and nuclear DNA sequences obtained from two museum specimens of *Lophichthys boschmai*. Employing ancient DNA techniques, we successfully extracted and analyzed the degraded and fragmented DNA. Our findings, derived from both mitochondrial genomes and nuclear ultra-conserved elements (UCE), reveal the Lophichthyidae as sister lineage to Tathicarpidae within the Antennarioidei. This study showcases the use of whole-genome “shotgun” sequencing data from formalin-fixed museum specimens in the context of an available UCE dataset. Given the enhanced versatility in data reusability this approach offers, we suggest it should be a favored approach when studying rare specimens.

## INTRODUCTION

Genetic data have long proven invaluable for modern taxonomic research (Hebert and Gregory, 2005). Molecular sequences have facilitated species identification (Waugh, 2007), revealed cryptic species (Struck, et al. 2018), and resolved relationships among clades with convergent phenotypes (Kawahara, et al. 2008). One clade with particularly rich genetic resources are vertebrates (*e.g*., Yates et al., 2020). Yet, genetic data have so far been unavailable for three out of the 1,123 extant vertebrate families (Bánki et al., 2025). All of these three families are marine acanthomorph fishes, monotypic, and rarely sampled: Dentatherinidae, Hispidoberycidae, and Lophichthyidae.

The only representative of Dentatherinidae is *Dentatherina merceri*, Mercer’s tusked Silverside, a species that was described by Patten and Ivantsoff (1983) from a number of very small (up to 3 cm) specimens collected in the Indopacific. It was initially placed within the Old World silversides family Atherinidae, but considered the sister taxon to Phallostethidae by Parenti (1984) on the basis of three shared derived characters. Parenti thus raised Dentatherininae to family level, a rank that has since been generally accepted (Ivantsoff et al., 1987; Campanella et al., 2015; Nelson et al., 2016; Bánki et al., 2025; but see Near and Thacker, 2024).

The family Hispidoberycidae is represented by the Bristlyskin, *Hispidoberyx ambagiosus*, a deep-water species of up to 18 cm length that is known from only a handful of specimens (Kotlyar, 2004). The first two of these (ZMMU 15416 and uncatalogued) were collected in bottom trawls in the southeastern Indian Ocean in 1979 (Kotlyar, 1981), and few further specimens were added from the South China Sea in subsequent years (Yang et al., 1988; Kotlyar, 1991). While the family was erected to accommodate the character combination of these specimens, which was seen as intermediate between Berycoidei and Stephanoberycoidei (Kotlyar, 1981), a phylogenetic analysis based on morphological characters suggested a position as sister taxon to the family Stephanoberycidae (Moore, 1993).

The sole member of Lophichthyidae is Boschma’s Frogfish, *Lophichthys boschmai*, a small (up to 7 cm) frogfish that is found in the Arafura and Timor Seas between Australia and New Guinea. The species and family were described from a single specimen (RMNH.PISC.24606) caught in 1954, (Boeseman, 1964), and few specimens have been collected afterwards (Hart et al., 2022). The family Lophichthyidae was erected by Boeseman because the character combination did not allow placement in any of the three anglerfish families considered at the time, Lophiidae, Antennariidae, and Chaunacidae. Nevertheless, Boeseman (1964) suggested an affinity with Lophiidae based on osteological similarities. In contrast, the more extensive osteological comparison by Pietsch (1981) suggested a position within a clade formed by Antennariidae, Tetrabrachiidae, and Brachionichthyidae, supported by two synapomorphies. While phylogenomic analyses have since identified Antennariidae as a polyphyletic taxon, its monophyletic components were found to form a well-supported clade – the suborder Antennarioidei – together with Tetrabrachiidae and Brachionichthyidae (Hart et al., 2022; Miller et al., 2024). Due to its lack of genetic data, the family Lophichthyidae could not be included in this phylogenomic analysis, but was nevertheless expected to fall within Antennarioidei based on shared morphological features and geographic distribution (Hart et al., 2022).

A fourth acanthomorph family, Scytalinidae, lacked genetic data until recently, but sequences of ultra-conserved elements (UCEs) have now been produced for its only representative, the Graveldiver *Scytalina cerdale*. The phylogenetic study enabled by this dataset revealed a phylogenetic position of *Scytalina* clearly nested within Stichaeidae (Ghezelayagh et al., 2022).

For rare and elusive species such as *Dentatherina merceri*, *Hispidoberyx ambagiosus*, and *Lophichthys boschmai*, fresh tissue samples suitable for DNA sequencing are difficult to obtain. Most of the few available specimens of these species were collected decades ago and preserved without consideration for molecular genetic analyses. To better retain the specimens’ original shape, they were usually fixed in formalin, a solution of formaldehyde that hardens biological tissues by forming cross-links between proteins (French and Edsall, 1945). Unfortunately, however, formalin also cross-links DNA covalently to proteins, making it much less accessible (Do & Dobrovic, 2015).

Formalin-fixed museum specimens have therefore long been largely inaccessible for genetic studies. However, this situation has begun to change, as new “museomics” laboratory protocols have emerged, allowing the extraction of fragmented DNA from formalin-fixed specimens (*e.g.*, Campos & Gilbert,

2012; Straube et al., 2021; Hahn et al., 2024). When combined with highly efficient library preparation methods, which were originally developed for analysis of minute amounts of degraded ancient DNA (Gansauge and Meyer, 2013; Gansauge et al., 2017), it now allows the retrieval of usable genetic information from many museum specimens (Card et al., 2021; Straube et al., 2021; Raxworthy and Smith, 2021; Hahn et al., 2022).

Here, we apply techniques developed for the extraction and analysis of degraded, fragmented DNA – such as that from museum specimens – to fill one of the three remaining gaps in sequence availability for vertebrate families. We extract and sequence mitochondrial and nuclear DNA from two specimens of *Lophichthys boschmai*, the only representative of Lophichthyidae, which were collected in 1990 and 1996, likely both fixed in formalin, and stored at the Museum and Art Gallery of the Northern Territory (MAGNT) Ichthyological Collection. We succeed in reconstructing the entire mitochondrial genome of both specimens, alongside sufficient nuclear sequence information to include the species in a phylogenomic analysis based on UCEs. This analysis allows us to unambiguously identify the position of Lophichthyidae as the sister taxon to Tathicarpidae, and to estimate the divergence time between these two families at around 10.2 million years ago.

## MATERIALS & METHODS

### Sampling

We obtained material of two *Lophichthys boschmai* specimens from the Museum and Art Gallery of the Northern Territory (MAGNT), Australia: One whole specimen (catalogue number S.14353-006; Fig. 1) and a tissue plug and a gill arch of a second (S.14545-001). Both of these had been collected in the Joseph Bonaparte Gulf, an inlet of the Timor Sea west of Darwin, Australia. Specimen S.14353-006 was collected in 1996 at a depth of ∼22 m at coordinates 14.52°S, 128.87°E while S.14545-001 was collected in 1990 at a depth of ∼60 m at 14.04°S, 128.05°E.

**Figure 1:**
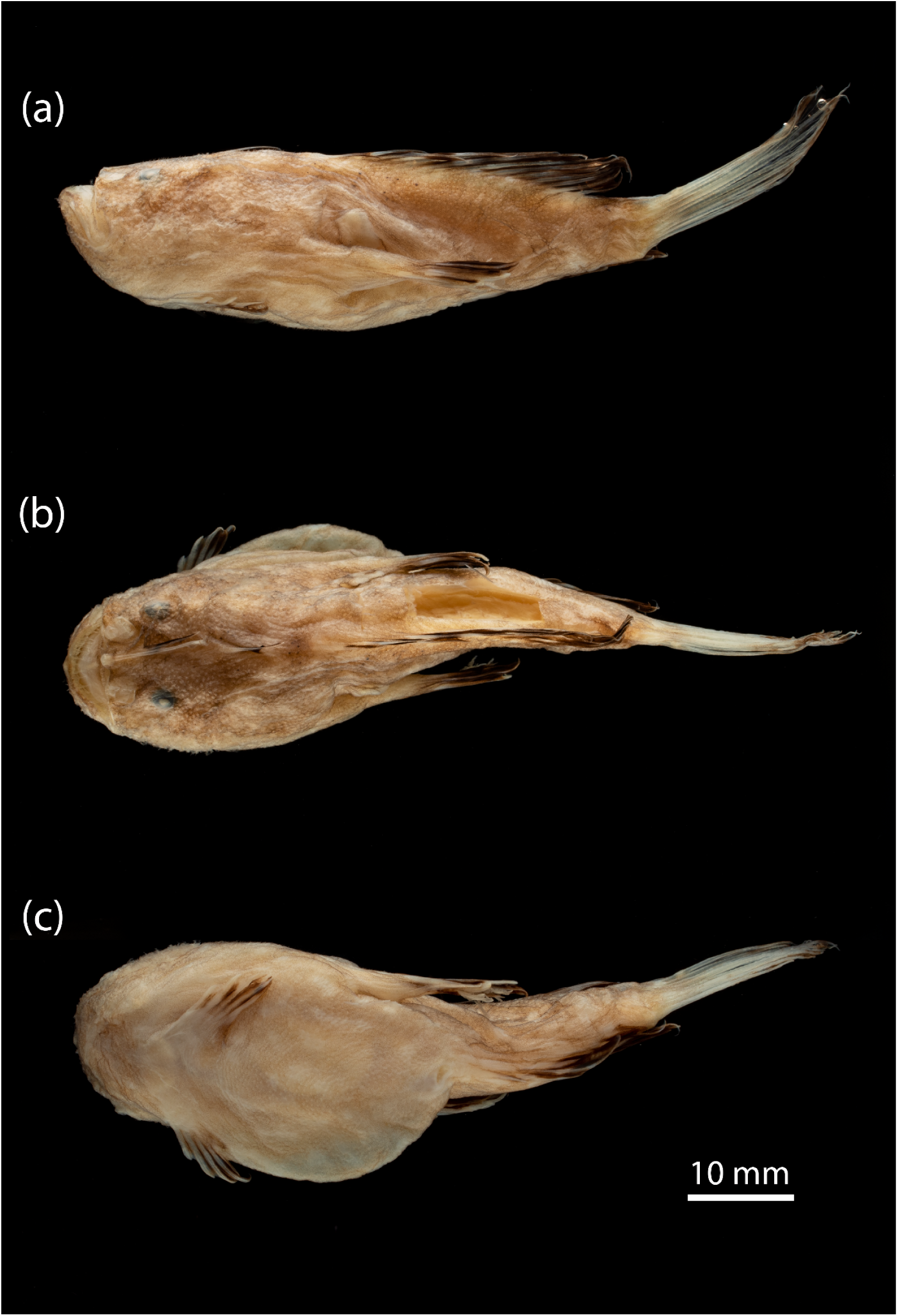
Photographs of specimen MAGNT S.14353-006. (a) lateral, (b) dorsal, and (c) ventral aspect.

Samples had likely both been treated with formalin solution and then stored in ethanol, but for sample S.14545-001 this can not be verified from records. The white pupils exhibited by specimen S.14353-006 (Fig. 1) – a feature often taken to indicate direct preservation in ethanol – may result from freezing prior to fixation with formalin (M. Hammer, pers. comm.). Residual formaldehyde concentration in the preservation fluid of specimen S.14353-006 was measured semi-quantitatively using MQuant colorimetric formaldehyde test strips (Merck) and was shown to be below the limit of detection (<10 mg/L).

### DNA extraction and sequencing

All experimental steps were performed as described in Muschick et al. (2022), which reports on other data produced in the same batch of reactions. Sensitive experimental steps prior to library amplification were performed in a dedicated laboratory to reduce the risk of contaminating the sample with exogenous DNA. We used the entire sample of S.14545-001 (14.1 mg dry weight) and a piece of muscle tissue from the right flank of S.14353-006 (31.1 mg dry weight) for DNA extraction. Dried tissue samples were immersed in 300 µL lysis buffer (260 μL ATL buffer (Qiagen) and 40 μL Proteinase K [20 mg/mL]). Reactions were incubated for 24 hours at 56°C. After centrifugation at 17,000×*g* for 5 min, 300 μL of the lysates’ supernatant were mixed with 3000 μL Buffer PB (Qiagen) and loaded onto MinElute spin columns (Qiagen) by repeated loading of 600 µL and centrifugation at 8000×*g* for 1 minute. Columns were washed twice with 600 μL Buffer PE (Qiagen) and dried by centrifugation for 1 min at 16,000×*g*. DNA was eluted into 50 μL of Buffer AE (Qiagen). Samples S.14545-001 and S.14353-006 yielded concentrations of 2.54 and 2.65 ng/μL, respectively. Two extraction negatives, further described in Muschick et al. (2022), contained 0.0244 ng/μL or were below the detection threshold of 0.02 ng/μL. Twenty and 19 μL of samples’ extracts, equivalent to 50.8 and 50.35 ng of DNA, respectively, were then used to build Illumina sequencing libraries by single-stranded DNA library preparation as described by Gansauge et al. (2017, 2020), using reagents as described in Gansauge et al. (2020). Libraries were uniquely dual indexed (7-bp index length) (Kircher et al., 2012), amplified until the end of the exponential phase and purified using the MinElute PCR purification kit (Qiagen). Libraries were pooled and size-selected to the range of 160 to 250 bp on a Blue Pippin instrument using a 3% cassette with internal markers. Both libraries were sequenced as part of a larger pool on an Illumina NextSeq500 instrument using a high-output single-end 75-bp read length kit with custom primers for read 1 (CL72) and index 2 (“Gesaffelstein”; Paijmans et al., 2017). Raw read files were demultiplexed using bcl2fastq v.2.19.1 (https://support.illumina.com/sequencing/sequencing_software/bcl2fastq-conversion-software.html), allowing for a maximum combined distance of 1 between barcodes, and saved to fastq format. In the same step, prevalent adapter sequences were trimmed from the ends of reads and reads shorter than 20 bp of length were omitted.

### Sequence analysis

To reconstruct the mitochondrial genomes of both *Lophichthys* samples we used an iterative mapping strategy. First, reads from both samples were joined and mapped iteratively against four Lophiiformes reference genomes which were padded with 200 “N”s at their start and end. The mapping proceeded as described in Muschick et al. (2023) by changing the reference sequence at called sites to produce a new, chimeric reference sequence for the next iteration. References used for this initial mapping were *Histrio histrio* (OP035243.1), *Dibranchus* sp. (MW080645.1), *Oneirodes thompsoni* (NC_013871.1) and *Chaunax pictus* (AB282833.1). For each initial reference, the iterative mapping proceeded until no additional reads could be mapped. Resulting called sequences were aligned with MAFFT v.7.470 and a majority-rules consensus calculated in R. Start and end of the linearised sequence (*i.e.*, the tRNA-Phe/D-loop transition in the circular sequence) were reconstructed by iteratively extending each end of the consensus into degenerate sequence (“N”) until the overlap could be identified. Reads of individual samples were then mapped to this completed consensus, which revealed a stretch with misaligning reads in the D-loop in one of the samples, caused by repeat unit variation. References were manually adjusted by varying the number of this sequence repeat to achieve a conflict-free mapping with even coverage depth for each sample. To assess the advantage of iterative mapping over single-pass mapping to reconstruct mitochondrial genomes from short reads with post-mortem damage and only distantly related references being available, we mapped reads from either specimen to the four references mentioned above, as well as the here reconstructed *Lophichthys* mitochondrial genomes, using the PALEOMIX pipeline (Schubert et al. 2014).

To better resolve its phylogenetic context, we also reconstructed additional mitochondrial genomes of putative close relatives of *Lophichthys* from available short read data, again employing an iterative mapping strategy. The *Tetrabrachium ocellatum* mitochondrial genome was reconstructed from read data first published in Swann et al. (2020), using the published mitochondrial genome from Miya et al. (2010) as starting reference. The two resulting large sequence blocks were then padded with degenerate sequence and iteratively mapped individually until the sequences could be merged, revealing the gene order already suggested by Fonseca et al. (2014). Mapping again to this sequence gave conflict-free, complete coverage, except for the identical parts in two control regions. The mitochondrial genome of *Tathicarpus butleri* was reconstructed for the first time using data published in Hart et al. (2022). The paired reads of 2×150 bp length in Hart et al.’s (2022) data were first deconstructed into stretches of 50 bp and added to the original data to aid the iterative mapping in proceeding over less conserved sequence. These duplicated reads were later ignored when calculating the average depth of coverage. We used our new *Lophichthys* mitochondrial genome sequence as starting reference and reconstructed almost the entire mitochondrial genome, lacking only parts of the second copy of a duplicated control region. All four newly reconstructed mitochondrial genome sequences were annotated using MitoFish v.3.98 (Zhu et al., 2023).

To verify that the reconstructed sequences for the two *Lophichthys boschmai* specimens were not affected by reference bias, we complemented the mapping-based sequence analysis with assembly of the mitochondrial reads. To do so, we applied the reference-independent sequence analysis workflow as described in Muschick et al. (2022). In brief, we used aTRAM v.2.0 (Allen et al., 2018) to locally assemble reads mapping to query sequences in nucleotide and amino acid format. As queries, we used all genes (protein coding, rRNA, or tRNA) of all 67 mitochondrial genomes for Lophiiformes that were available on NCBI on 4 December 2021. The contigs produced by aTRAM were further assembled with MIRA v.4.9.6. To compare these assembly-based sequences to the mapping-based mitochondrial genome sequences produced for the two *Lophichthys boschmai* specimens, we aligned these to each other with MAFFT v.7.470 (Katoh and Standley, 2013).

For complementary phylogenetic analyses based on nuclear data, we used the concatenated 75% complete data matrix alignment of UCE sequence data which underlies the phylogenomic trees of Hart et al. (2022). Because this matrix was produced from piled-up sequences from a hybridisation sequence capture experiment, it could not readily be merged with our data. Instead, we filtered and manipulated the concatenated UCE sequences of three individuals (Brachionichthys_sp_B1, Tathicarpus_butleri_A49, Tetrabrachium_ocellatum_C) and used them for mapping of the joined reads from specimens S.14353-006 and S.14545-001. Filtering and manipulation of the gappy UCE sequences extracted from the alignment of Hart et al. proceeded by first replacing stretches of four or more gap (“-”) or unknown (“?”) characters with “N”, then retaining only continuous or near-continuous (gaps of size 3 bp or smaller ignored) defined sequence stretches of 100 or more bp, while converting any shorter stretches to “N”. Subsequently, retained gaps of size 3 bp or smaller were deleted from each sequence. This produced degenerate sequences of ∼1.3 Mbp length, interspersed with stretches of 100 bp or longer defined sequence (∼150 kbp in total). These manipulated, concatenated UCE sequences were then used for iterative mapping like described below for the mitochondrial references. The three called sequences resulting from mapping the joined *Lophichthys* reads against three UCE references were then used to produce a consensus by deleting all “N”, aligning the remainders using MAFFT v.7.470, and retaining only sites with identical calls for all references. The resulting sequence was aligned to the concatenated 75% complete data matrix from Hart et al. (2022), downsampled to individuals given in Supplementary Table S1, using MAFFT.

### Phylogenetic analyses

Phylogenetic analyses were conducted separately for the mitochondrial genome and for nuclear genome-wide UCE loci. For phylogenetic analysis of the mitochondrial genome, we retrieved complete mitochondrial genome sequences of 49 acanthomorph fish species from NCBI, including 45 species of lophiiforms, and, as outgroups, the caproiform *Antigonia capros* and three species of the genus *Chaetodon*. From each of these 49 mitochondrial genomes and the four newly generated mitochondrial genomes for *Lophichthys boschmai, Tathicarpus butleri* and *Tetrabrachium ocellatum*, we extracted the sequences for the 12S and 16S rRNA genes as well as all 13 protein-coding genes of the mitochondrial genome. This was done through BLASTN and TBLASTN (Altschul et al., 1990) searches, with nucleotide and amino-acid query sequences selected from the *Antigonia capros* mitochondrial genome (NCBI accession number AP002943.1; Miya et al., 2001). Per marker, sequences were aligned using MAFFT v.7.505, and the resulting alignments were subsequently concatenated while storing information for four partitions: One each for all first, second, and third codon positions (CP), and one for the concatenated rRNA genes. The first three of these partitions each contained 3,768 bp and the fourth had 2,834 bp, for a total alignment length of 14,138 bp. This concatenated and partitioned alignment was then used as input for IQ-TREE v.2.2.2 (Minh et al., 2020) to reconstruct the mitochondrial phylogeny under maximum likelihood. We employed IQ-TREE’s inbuilt standard model selection (Kalyaanamoorthy et al., 2017) followed by tree inference. Supported by the Bayesian information criterion (Schwarz, 1978), derivatives of the general time-reversible (GTR) model (Tavaré, 1986) were selected for the three CP-specific partitions (CP1: GTR+F+I+R4; CP2: GTR+F+R4; CP3: GTR+F+I+R5) and the TIM2+F+I+R4 model was selected for the rRNA partition. Node support was assessed with 1,000 ultrafast bootstrap (BS) replicates (Hoang et al., 2018).

For phylogenetic analysis of nuclear UCE sequences, we concatenated all individual UCE alignments into a single large alignment with a length of 1,000,732 bp. The maximum-likelihood phylogeny for this alignment was then reconstructed with IQ-TREE2, again using the inbuilt standard model selection, in which the Bayesian information criterion supported the TVM+F+I+R5 substitution model. Node support was again assessed with 1,000 ultrafast bootstrap replicates.

Finally, we also used the nuclear UCE sequence alignment for Bayesian inference of the time-calibrated phylogeny with the program BEAST2 (Bouckaert et al., 2019). For feasible run times of the computationally demanding Bayesian analyses, however, we first filtered the UCE alignment with BMGE v.1.0 (Criscuolo and Gribaldo, 2010), enforcing a maximum proportion of missing data per site of either 0.02 (“strict” filtering) or 0.05 (“permissive” filtering) and a maximum entropy score of 0.7. The resulting filtered alignments had lengths of 38,815 and 93,313 bp, respectively. Both alignment files were checked visually for obviously misaligned sequences, which were then replaced with missing data. Input files for BEAST2 were generated with the R library babette (Bilderbeek and Etienne, 2018). We applied a relaxed clock model (Drummond et al., 2006) and a birth-death model as tree prior (Gernhard, 2008). The clock model was calibrated through an age constraint on the divergence of Lophiidae and other lophiiforms, matching the divergence time between *Lophius vaillanti* and *Antennarius striatus* estimated by Musilova et al. (2019). This age constraint was specified as a lognormal prior distribution with a mean of 63.99 Ma and a standard deviation of 0.15. We applied the general time-reversible (GTR) model of sequence evolution (Tavaré, 1986) with gamma-distributed rate variation in four rate categories. We ran five replicate BEAST2 analyses, each with 75 million Markov-chain Monte Carlo (MCMC) iterations. Convergence and stationarity of the MCMC were assessed by inspection of trace plots in Tracer v.1.7 (Rambaut et al., 2018) and confirmed through effective sampling sizes (ESS) greater than 200. The posterior tree distributions were summarized in the form of a maximum-clade-credibility tree generated with TreeAnnotator v.2.6.7 (Heled and Bouckaert, 2013).

## RESULTS

### Sequencing and reconstruction of *Lophichthys boschmai*’s mitochondrial genome

Pooled sequencing resulted in 19.6 and 56.5 million reads for specimens S.14353-006 and S.14545-001, respectively. Iterative mapping of joined reads of both individuals to mitochondrial references proceeded for up to 48 iterations (to *Chaunax* sp.) and an additional 3 iterations for mapping individual specimens’ reads to the consensus of the mapping of joint reads to four Antennarioidei references. In their final iteration, specimens S.14353-006 and S.14545-001 produced 14,932 and 28,961 unique hits, with average read lengths of 43.2 and 54.5 bp and average depth of coverage of 38.5 and 94.6, respectively. The complete mitochondrial genome sequences reconstructed from iterative mapping are 16,610 and 16,647 bp in length. Mapping the reads iteratively proved necessary, as single-pass mapping to any of the four reference mitochondrial genome sequences produced only 1,314 unique hits for both specimens combined in the best case (to reference *Histrio histrio*). Additionally, coverage from single-pass mapping was also uneven, as may be expected when using a phylogenetically distant reference. While conserved regions in parts of the 16S ribosomal RNA gene attracted numerous hits, most of the mitochondrial genome (78.5%) received no coverage. The same approach using the final, reconstructed sequence from *Lophichthys boschmai* specimen S.14545-001 as reference yielded a combined 43,748 unique hits – representing a 33.3-fold increase over single-pass mapping to the phylogenetically closest reference available.

Our assembly-based reconstruction of the mitochondrial genome produced five contigs with a total length of 8,784 bp for specimen S.14353-006 and two contigs with a total length of 13,662 bp for S.14545-001. All of these contigs mapped uniquely to the mitochondrial genome sequences produced through iterative mapping, with only three single-nucleotide differences (0.01% of nucleotides; Fig. 2, Supplementary Table S2). All of these differences were near the ends of a contig (one for S.14353-006 and two for S.14353-006) and are therefore considered likely assembly errors.

**Figure 2:**
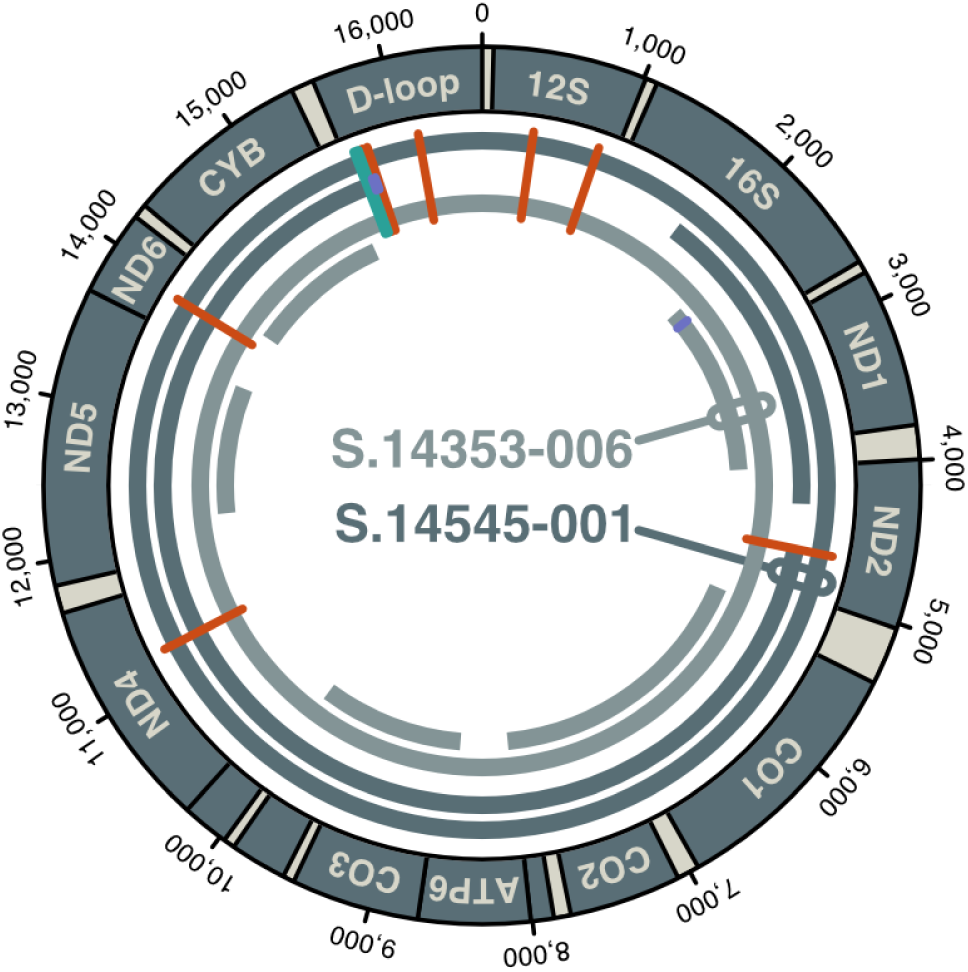
The mitochondrial genome of *Lophichthys boschmai*. Coordinates are in units of base pairs. The four complete and partial circles in dark gray and light gray represent the mitochondrial genome sequences based on mapping (the complete outer circle of each pair) or assembly (the fragmented inner circle of each pair). Light gray circles correspond to sequences of S.14353-006, while dark gray circles represent sequences of S.14545-001. Eight single-nucleotide differences between these sequences are highlighted in red; one 37-bp indel in the D-loop region is shown in green, and three putative single-nucleotide errors at the ends of two assembled fragments are marked in purple (see Supplementary Table S2).

The comparison of the two complete mitochondrial genome sequences based on iterative mapping showed close relatedness of the two *Lophichthys boschmai* specimens. Excluding the three probable assembly errors, the two sequences were separated by no more than eight single-nucleotide differences in addition to one indel of 37 bp. The indel and three single-nucleotide differences were found in the D-loop region; two single-nucleotide differences are located in the 12S rRNA gene, and the remaining three are in the protein-coding genes ND2, ND4, and ND6. Of the latter three, two are non-synonymous while the third substitutes a Valine for a Methionine in ND6 (Fig. 2).

### Reconstruction of mitochondrial genomes of *Tathicarpus butleri* and *Tetrabrachium ocellatum*

Available raw sequencing data, presumably collected with different objectives than reconstructing the mitochondrial genome, nevertheless contained sufficiently many unique mitochondrial reads to assemble all coding and most non-coding parts of both species’ mitogenomes. The finally achieved average depth of coverage was 74 for *Tathicarpus butleri* and 35 for *Tetrabrachium ocellatum*. While the quite distinct gene order in *Tetrabrachium* had been remarked on before (Fonseca et al., 2014), for *Tathicarpus*, we here discover a rearrangement of the ND6 and tRNA-Glu genes to a position in between the copies of a duplicated control region. Despite the wide divergence from the references used as starting points for iterative mapping (*i.e*., *Histrio histrio*, *Dibranchus* sp., *Oneirodes thompsoni* and *Chaunax pictus* in case of *Lophichthys boschmai,* and *Lophichthys boschmai* in case of *Tathicarpus butleri*), it was possible to reconstruct all or almost all of the targeted mitochondrial genomes, lacking only sequence-identical parts of duplicated control regions in *Tathicarpus butleri*.

While the use of *Lophichthys boschmai* as a reference for the reconstruction of the *Tathicarpus butleri* mitogenome could be expected to cause a reference bias that artificially reduces the divergence between the two species, this is unlikely to have taken place here. The final alignments of all mitochondrial genomes reconstructed by iterative mapping had deep, even coverage and are free of heterozygous positions, indicating strong internal consistency of the data.

### Post-mortem damage and taxonomic assignment

As expected in decades-old, formalin-fixed, ethanol-preserved tissue, a pattern of elevated deamination damage towards the ends of DNA fragments is observed, albeit at a relatively low level (Supplementary Figure S1) when compared to similarly preserved museum specimens (Muschick et al., 2022) or sediment-embedded fish subfossils (Muschick et al., 2023). Mapping to NCBI’s non-redundant sequence database showed that a large proportion of reads from both *Lophichthys boschmai* specimens were assigned to percomorph taxa, indicating their endogenous origin (Supplementary Figures S2 and S3).

Partitioned maximum-likelihood phylogenetic analysis of the two mitochondrial genome sequences for *Lophichthys boschmai* with IQ-TREE, alongside those of 52 other percomorph species, unambiguously supported the monophyly of the two *Lophichthys boschmai* specimens (BS 100%), which were connected with a near-zero branch length. The mitochondrial phylogeny reproduced lophiiform relationships established with genome-wide markers (Hart et al., 2022; Miller et al., 2024): The included representatives of Lophioidei were placed as the sister group to all other lophiiform taxa (BS 95%); Chaunacoidei formed a monophyletic group with Ceratioidei (BS 100%); and Ogcocephaloidei were supported as the sister group to Antennarioidei, albeit weakly (BS 58%). As expected from morphological data (Pietsch, 1981), *Lophichthys boschmai* was placed within Antennarioidei (BS 100%), where the species clustered with the representative of Tathicarpidae, *Tathicarpus butleri* (BS 100%; Fig. 3).

**Figure 3:**
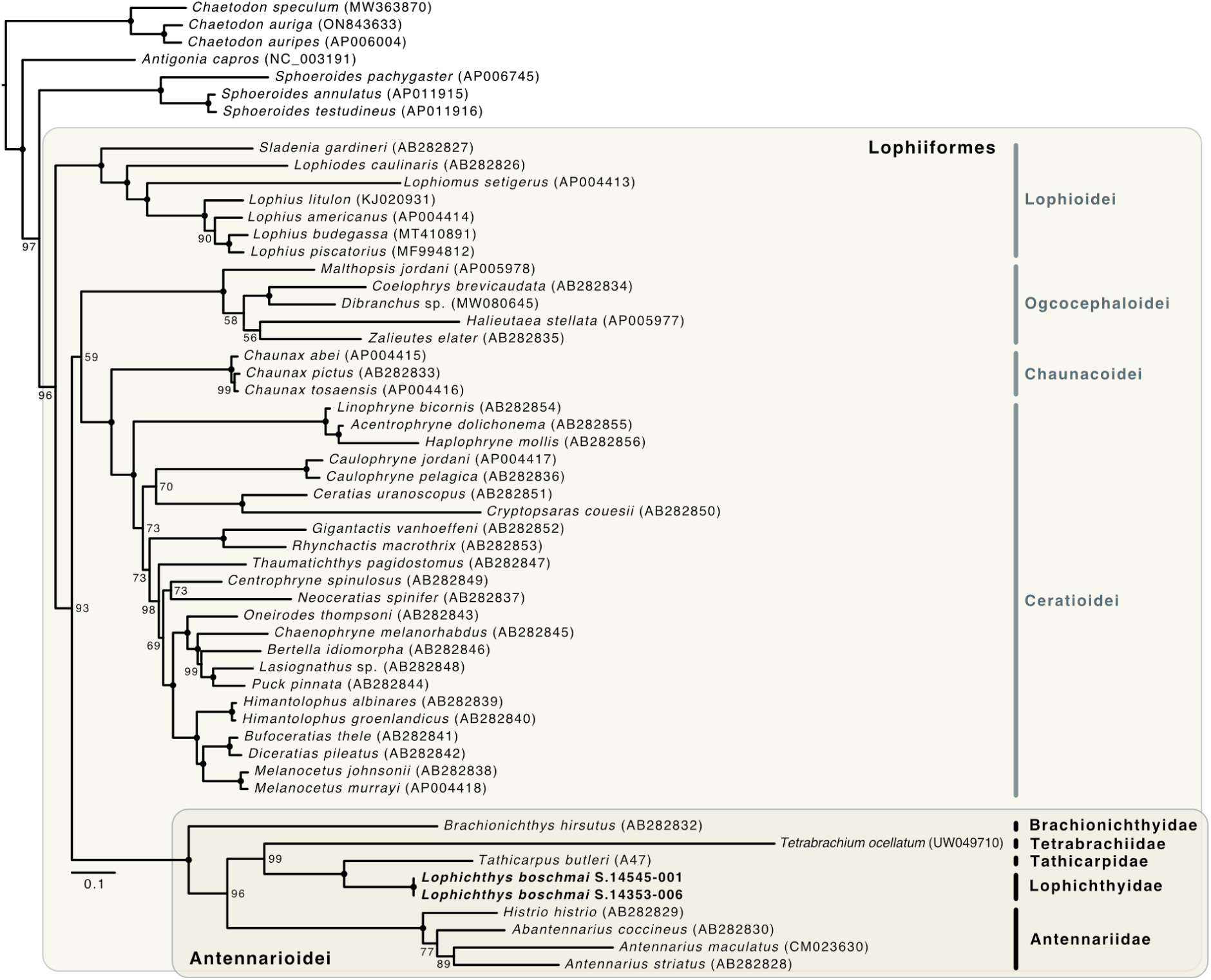
Maximum-likelihood phylogeny reconstructed from mitochondrial genome sequences. The phylogeny was reconstructed with IQ-TREE. Black circles on nodes indicate full (100%) bootstrap support; labels mark nodes with bootstrap support below 100%. The order Lophiiformes, lophiiform suborders, and families within the suborder Antennarioidei are highlighted.

Applying maximum-likelihood phylogenetic analysis to concatenated alignments of genome-wide UCE sequences for 51 species corroborated the results of the mitochondrial phylogeny. Again, the same relationships among the five lophiiform suborders were recovered, in each case with full node support (BS 100%). Within Antennarioidei, the UCE dataset included sequences of 32 species and thus allowed a greater resolution of the suborder’s internal relationships than the mitochondrial dataset. Unsurprisingly given the overlap in sequence data, we recovered nearly the same topology of Antennariidae, Tathicarpidae, Histiophrynidae, Brachionichthyidae, and Rhycheridae as Hart et al. (2022) and Miller et al. (2024) (Fig. 4). In this topology, Antennariidae are the sister group to a clade formed by all other families (BS 100%), Brachionichthyidae cluster with Rhycheridae (BS 100%), and Tetrabrachiidae group with Tathicarpidae and Histiophrynidae (BS 81%). However, in contrast to Hart et al. (2022) and Miller et al. (2024), our analysis did not place Tetrabrachiidae with Tathicarpidae, but found the latter to be more closely related to Histiophrynidae (BS 99%).

**Figure 4:**
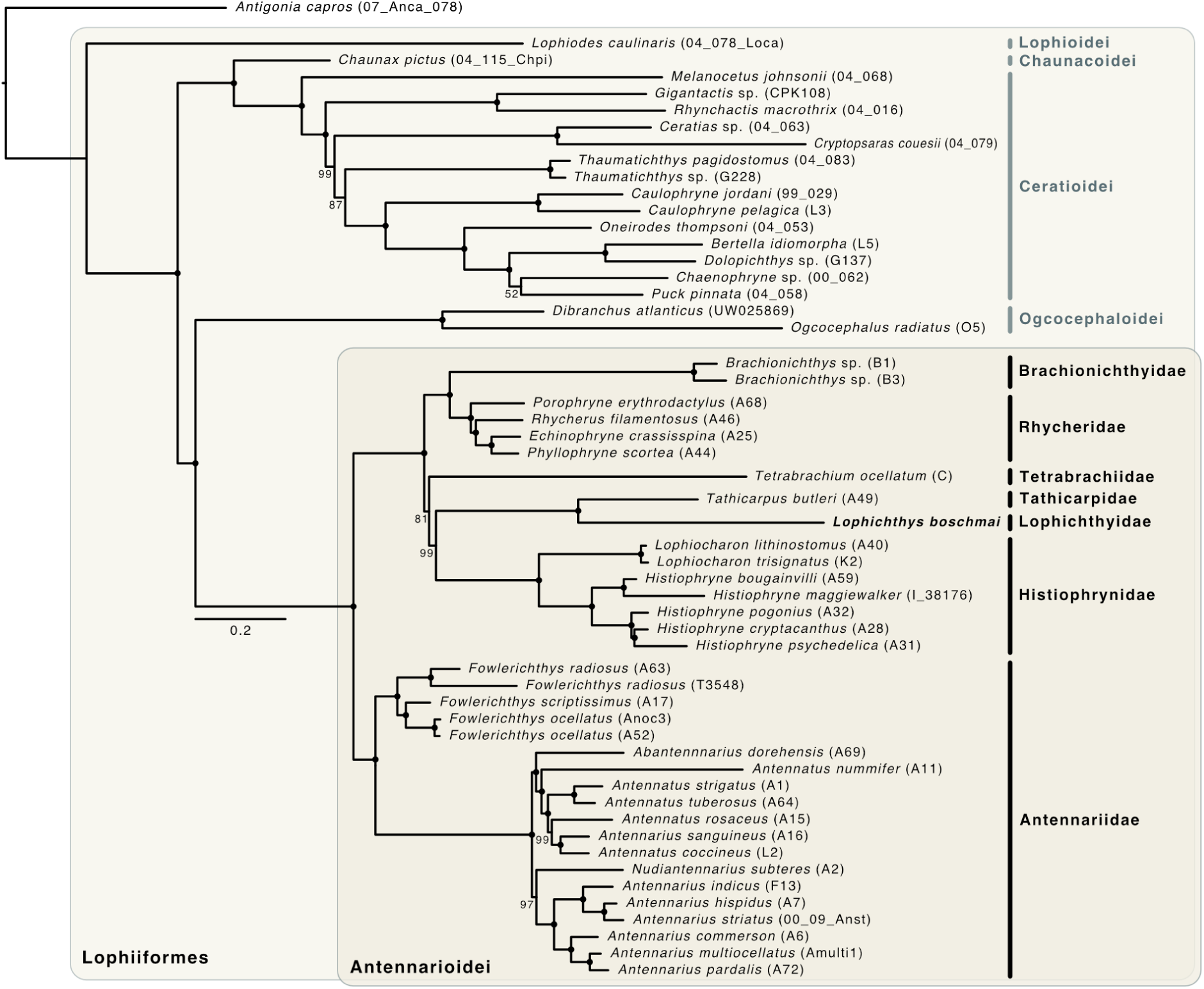
Maximum-likelihood phylogeny reconstructed from genome-wide UCE marker sequences. The phylogeny was reconstructed with IQ-TREE. Black circles on nodes indicate full (100%) bootstrap support; labels mark nodes with bootstrap support below 100%. The order Lophiiformes, lophiiform suborders, and families within the suborder Antennarioidei are highlighted.

*Lophichthys boschmai* was, with full support (BS 100%), placed within this group, as the sister group of *Tathicarpus butleri*, the sole member of Tathicarpidae. This position does not conflict with the mitochondrial phylogeny, as Histiophrynidae were not included in the mitochondrial dataset.

However, the relative positions of Brachionichthyidae and Antennariidae were discordant between the two phylogenies, as Antennariidae appeared as the sister group to all others in the UCE-based phylogeny (BS 100%; Fig. 4) while Brachionichthyidae took that position in the mitochondrial phylogeny (BS 99%; Fig. 3).

Complementing the maximum-likelihood phylogenetic analyses, we also performed Bayesian inference of the phylogeny based on the UCE dataset. Allowing a maximum of two missing nucleotides per site (equivalent to a maximum proportion of missing data of 0.05), the resulting time-calibrated phylogeny recovered a strongly supported topology similar to that of the maximum-likelihood inference from the same dataset (Fig. 5). Quantified as Bayesian posterior probability (BPP), all but three of the nodes in this topology received full bootstrap support (BPP 1.0). Lophioidei once again were the sister group to a clade formed by two reciprocally monophyletic pairs of suborders, combining Chaunacoidei and Ceratioidei on the one hand and Ogcocephaloidei and Antennarioidei on the other. Within Antennarioidei, the family Antennariidae was again in a sister-group position to a clade formed by all other families; Brachionichthyidae and Rhycheridae clustered with each other; and so did Tathicarpidae and Lophichthyidae, all with full support (BPP 1.0). However, unlike in the maximum-likelihood phylogeny, the latter two families appeared closer to Tetrabrachiidae than to Histiophrynidae (BPP 0.88). The inferred timeline suggests that extant lineages of Antennarioidei began to diversify in the Oligocene, around 27.0 million years ago (Ma) with a 95% highest-posterior-density (HPD) interval ranging from 38.7 to 16.4 Ma. The divergence of Lophichthyidae from its unambiguous sister group, Tathicarpidae, most likely took place in the Miocene, around 10.2 Ma (95% HPD 17.3–4.1 Ma). Notably, this age between the two families is younger than several divergence events within other antennarioid families, such as the divergence of *Lophiocharon* and *Histiophryne* within Histiophrynidae (12.3 Ma; 95% HPD 19.0–6.1 Ma) or that of *Fowlerichthys* from *Antennarius* (and other genera) within Antennariidae (16.7 Ma; 95% HPD 24.9–9.4 Ma). The more strictly filtered version of the UCE dataset (allowing one missing nucleotide per site; enforced through a maximum proportion of missing data of 0.02) supported a phylogeny that was largely concordant, albeit with Tathicarpidae and Lophichthyidae placed as a weakly supported (BPP 0.57) sister group to Brachionichthyidae and Rhycheridae (Supplementary Figure 4).

**Figure 5:**
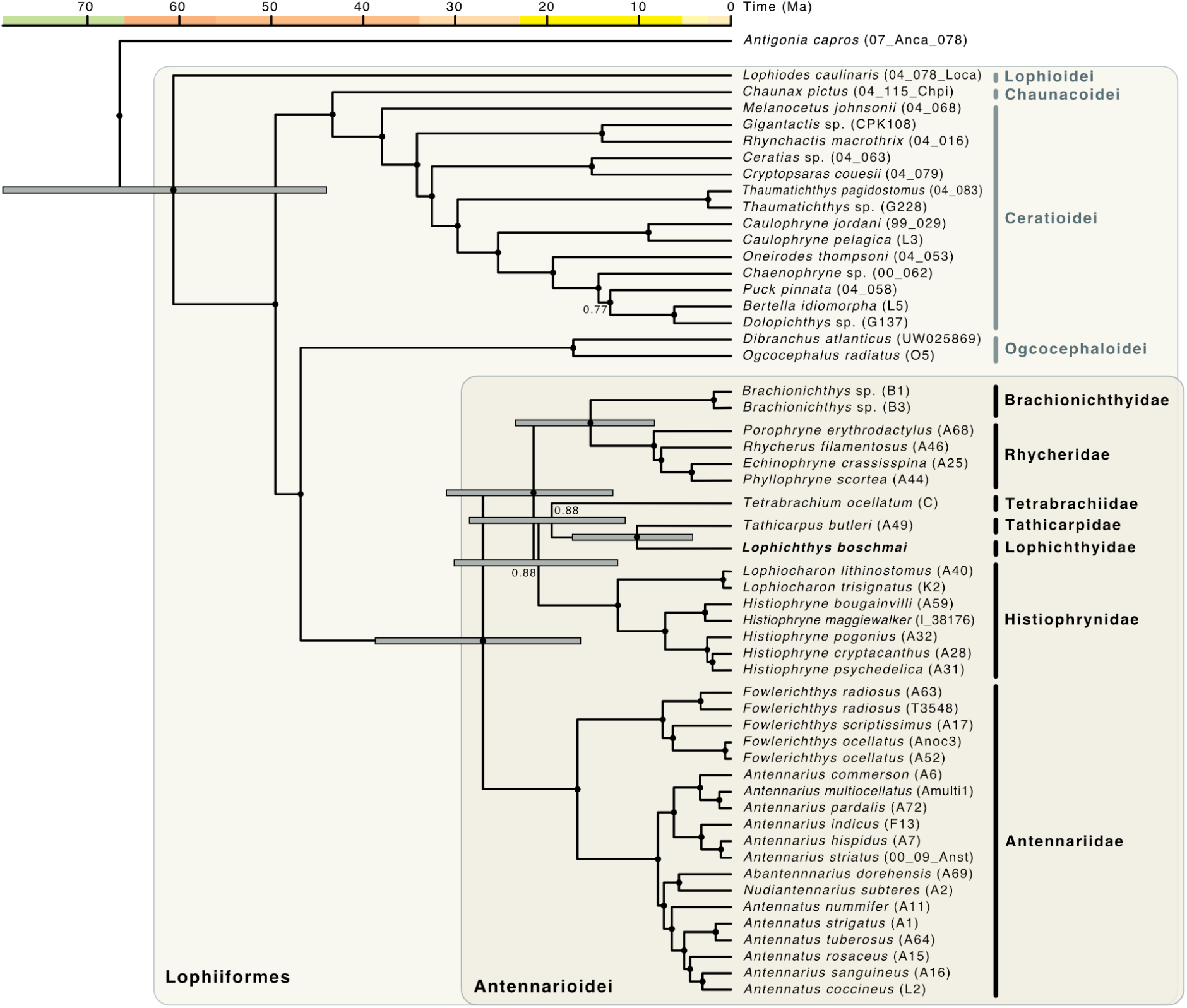
Time-calibrated phylogeny reconstructed from genome-wide UCE marker sequences, after permissive filtering for missing data (allowing a maximum of two missing nucleotides per site). The phylogeny was reconstructed by Bayesian inference with BEAST2. Black circles on nodes indicate full (1.0) Bayesian posterior probability; labels mark nodes with Bayesian posterior probabilities below 1.0. For the calibrated node (the first divergence among Lophiiformes) and for inter-familial divergences among Antennarioidei, node bars are shown to indicate 95% highest posterior density intervals for age estimates. The order Lophiiformes, lophiiform suborders, and families within the suborder Antennarioidei are highlighted.

Importantly, the placement of Lophichthyidae with Tathicarpidae received the same strong support (BPP 1.0) and their divergence time was not affected by the topological differences (10.5 Ma; 95% HPD 17.7–4.4 Ma).

## DISCUSSION

Our phylogenetic analyses based on mitochondrial and nuclear sequence data unambiguously placed *Lophichthys boschmai* – and thus the family Lophichthyidae – within the suborder Antennarioidei. This placement is according to expectation, given shared morphological characters with other antennarioid families, such as the loss of the mesopterygoid, a reduction or loss of the epural, and oval-shaped ovaries (Pietsch, 1981; Pietsch, 1984; Hart et al., 2022). More precisely, our mitochondrial and nuclear datasets all strongly support a position of Lophichthyidae as the sister group of Tathicarpiidae.

Bayesian inference of the time-calibrated phylogeny suggested a divergence of Lophichthyidae and Tathicarpidae in the Miocene (around 10.2 Ma, with a 95% HPD interval of 17.3–4.1 Ma). This age estimate is comparable to some within-genus divergence times (*e.g.,* that of *Caulophryne jordani* and *Caulophryne pelagica*, estimated at 9.0 Ma, with a 95% HPD interval of 15.8–3.4 Ma). It implies that Lophichthyidae and Tathicarpidae are the two youngest families within Antennarioidei, and possibly among all ray-finned fishes (Actinopterygii), a group with 488 families and 32,513 species (Bánki et al., 2025). In fact, the family-level, and largely complete, time-calibrated phylogeny of ray-finned fishes by Rabosky et al. (2018) includes not a single divergence younger than 20 Ma. While differences in the time calibration between our study and that of Rabosky et al. (2018) may prevent a direct comparison of these ages, it remains without doubt that Lophichthyidae and Tathicarpidae are among the youngest ray-finned fish families. This, together with the fact that both taxa are monotypic, casts doubt on whether their classification at the family level is warranted.

A counterargument justifying the family-level classification could be made if *Lophichthys boschmai* would be unusually distinct in its phenotype or genome. Genomically, this may indeed be the case, as the Bayesian phylogenetic analysis with BEAST2 assigned the highest inferred substitution rates in the entire phylogeny (*r* = 0.0018 substitutions per site per million years; 95% HPD 0.0007–0.0032) to the terminal branch leading to this *Lophichthys boschmai*. This high rate of molecular evolution is also reflected in the long terminal branch found in the maximum-likelihood phylogeny based on UCE data. However, given that our sequence data for *Lophichthys boschmai* derived from formalin-fixed museum specimens and the nuclear read coverage was comparatively low, an elevated frequency of erroneous base calls cannot be excluded as a potential cause of the apparently higher substitution rate.

The phenotypic rate of evolution of *Lophichthys boschmai* has so far not been quantified, and the species has overlapping character distributions with many other antennarioid taxa. It differs, for example, from *Antennarius* in the presence of a single row of epibranchial teeth and the absence of a pseudobranch and a swim bladder, but the latter two traits are shared with *Tetrabrachium* and *Brachionichthys* (Pietsch, 1981). In summary, given its young age, its monotypic nature, the ambiguity regarding its rate of molecular evolution, and the lack of strong phenotypic differences, the classification of Lophichthyidae at the rank of family may be questioned. This conclusion agrees with a recently proposed revision of ray-finned fish classification, which groups all members of Antennarioidei into the family Antennariidae (Near and Thacker, 2024).

According to our Bayesian analysis, Lophichthyidae, Tathicarpidae, and Tetrabrachiidae form a species-poor clade of four phylogenetically distinct species (including *Dibrachichthys melanurus* as a second member of Tetrabrachiidae; Pietsch et al. 2009) that began to diverge around 19.5 Ma (95% HPD 28.5–11.5 Ma). All of these four species are shallow-water frogfishes distributed along the northern coast of Australia in the Timor and Arafura Seas. They thus exemplify a trend recently identified for Lophiiformes, where paradoxically, species inhabiting the homogeneous bathypelagic realm have greater diversification rates than those of more heterogeneous coastal areas (Miller et al. 2024). This trend could partially be explained if frequent gene flow along the coast would prevent the formation of genetic differentiation within shallow-water species. This would also be consistent with the very low genetic distance between the mitochondrial genomes of the two sequenced *Lophichthys boschmai* specimens (9 genetic differences despite a geographic distance of over 100 km between their sampling sites). In marine fishes, such gene flow is commonly mediated by the dispersal of pelagic larvae with currents (Barlow, 1981). Pelagic rafting eggs are in fact found in frogfishes; however, this is not the case for *Lophichthys boschmai* and *Tetrabrachium ocellatum*, which both produce large eggs that hatch into demersal juveniles with limited dispersal potential (Pietsch et al., 2009).

Alternatively, diversification could be hampered by high rates of extinction in Lophichthyidae, Tathicarpidae, and Tetrabrachiidae. This scenario would support the long-standing hypothesis that non-dispersing taxa might be more prone to extinction due to random events extirpating local populations (Doherty et al., 1985). While the current distribution and habitat of the three families do not appear to be threatened (IUCN 2024), it is worth noting that most of their habitat lay dry when sea levels were lower during periods of Pleistocene glaciation (Hantoro et al., 1995). Thus, it appears plausible that habitat alterations in the past 20 million years disproportionately affected shallow-water frogfish taxa, reducing a diversity that may have been greater in the past.

This study demonstrates that moderate sequencing depth and shotgun sequencing can be sufficient to place museum specimens into a phylogenomic context. Many studies which generate and utilise sequences of UCE loci are performing target capture to enrich libraries in molecules of interest to make sequencing more economical. However, the resulting data which is then made available via repositories may lack in sequence reads from loci other than those targeted UCEs, and therefore be of limited value for re-use in other contexts. Especially in the case of rare museum specimens, possibly from extinct species, or type material, an opportunity is then lost to reduce the impact of destructive sampling to a single event when subsequent studies with different objectives have to produce new data from repeated sampling. Hence, for specimens of extraordinary value, the most sensitive methods should be chosen, such as single-stranded DNA library preparation, to maximise the yield from a minimally invasive sampling, and should be followed by a shotgun sequencing approach to ensure re-usability of the data.

With sequence data now being available for *Lophichthys boschmai*, only two vertebrate families remain without genetic information; Dentatherinidae and Hispidoberycidae. This means that the construction of the first complete family-level vertebrate phylogeny may soon be possible. Such phylogenies that are complete at a certain taxonomic rank are valuable not just from an ideological point of view, but also because they allow the assignment of all extant species diversity to their tips. This complete assignment is important, for example, for the accurate estimation of diversification rates, but also for the use of fossil constraints in time calibration (Matschiner et al., 2017). Thus, better-calibrated phylogenies for vertebrates may be on the horizon, and these may provide unprecedented information about vertebrate diversification through time and space.

## ACKNOWLEDGEMENTS

We thank Michael Hammer from the Museum and Art Gallery of the Northern Territory (MAGNT) for providing tissue samples of *Lophichthys boschmai* and Marcelo Sánchez from the University of Zurich for discussions as well as financial support. Computational analyses were performed on resources provided by Sigma2 – the National Infrastructure for High Performance Computing and Data Storage in Norway, and the Euler cluster of ETH Zurich. Access to laboratories at the ETH Zurich was kindly facilitated by Mark Lever. The Genetic Diversity Centre at ETH Zurich provided access to laboratory and HPC facilities. The Functional Genomics Centre Zürich provided assistance with sequencing.

## AUTHOR CONTRIBUTIONS

Moritz Muschick: Conceptualization, Formal Analysis, Investigation, Data Curation, Writing – Original Draft, Writing – Review & Editing, Visualization, Project Administration Lukas Rüber: Conceptualization, Resources, Writing – Review & Editing Michael Matschiner: Conceptualization, Formal Analysis, Resources, Data Curation, Writing – Original Draft, Writing – Review & Editing, Visualization, Project Administration

## DATA ACCESSIBILITY

Raw-read sequencing data for this study have been deposited in the European Nucleotide Archive (ENA) at EMBL-EBI under accession number PRJEB83588 (https://www.ebi.ac.uk/ena/browser/view/PRJEB83588). Mitochondrial genome sequences of both *Lophichthys boschmai* specimens are accessible in the NCBI Genbank repository under accession numbers PP328847 and PP328846. Mitochondrial genome sequences of *Tathicarpus butleri* and *Tetrabrachium ocellatum* are accessible on the Zenodo repository under DOI 10.5281/zenodo.14905955.

