## Supplementary Information for "Museomics reveals the phylogenetic position of an enigmatic vertebrate family (Lophiiformes: Lophichthyidae)"

Moritz Muschick

Lukas Rüber

Michael Matschiner

AUTHORS' AFFILIATIONS

M. Mu.: Aquatic Ecology and Evolution, Institute of Ecology and Evolution, University of Bern, Bern, Switzerland; Department of Fish Ecology and Evolution, EAWAG, Swiss Federal Institute for Aquatic Science and Technology, Kastanienbaum, Switzerland.
Current affiliation: Faculty of Biology, Ludwig-Maximilians-Universität München, Munich, Germany.

L. R.: Naturhistorisches Museum Bern, Bern, Switzerland; Department of Fish Ecology and Evolution, EAWAG, Swiss Federal Institute for Aquatic Science and Technology, Kastanienbaum, Switzerland.

M. Ma.: Natural History Museum Oslo, University of Oslo, Oslo, Norway; Department of Palaeontology and Museum, University of Zurich, Zurich, Switzerland.
Current affiliations: Faculty of Biology, Ludwig-Maximilians-Universität München, Munich, Germany; SNSB-Zoologische Staatssammlung München, Munich, Germany

SUPPLEMENTARY FIGURES

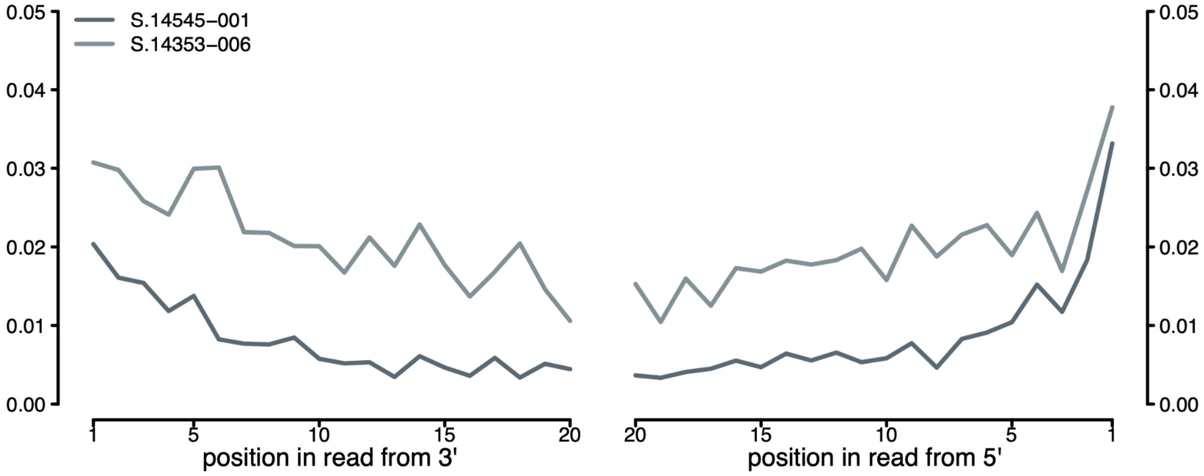


Supplementary Figure S1: Post-mortem damage in mapped mitochondrial reads. (a) Post-mortem damage from cytosine deamination is distributed unevenly along mapped reads. The fraction of T where the reference is C is plotted against the position in mapped reads either counted from the 3′-end or the 5′-end. As this chemical alteration is especially prevalent in single-stranded overhangs, the relative abundance of apparent C to T changes at reads' ends is indicative of damaged DNA.

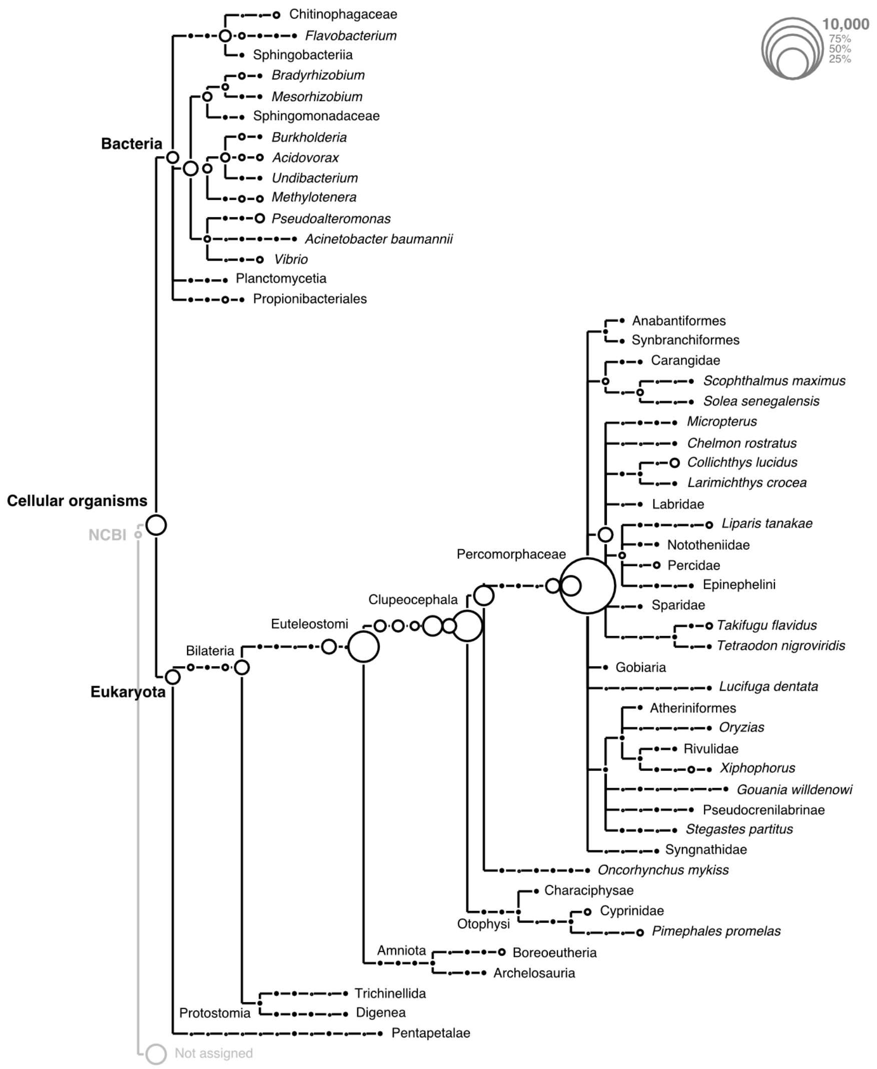


Supplementary Figure S2: Taxonomic assignment of individual reads for MAGNT S.14545-001. Reads were mapped to NCBI's non-redundant database with Diamond, and illustrated with MEGAN. Only nodes with a minimum support of 50 reads are shown.

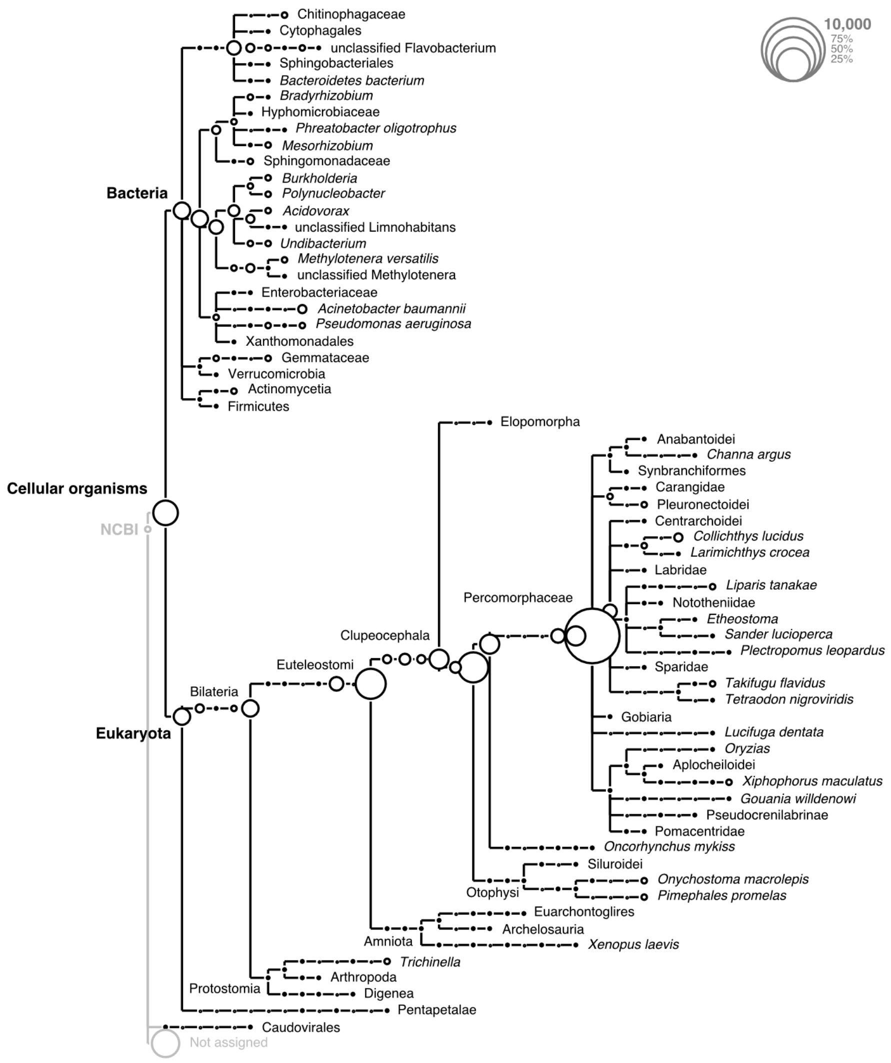


Supplementary Figure S3: Taxonomic assignment of individual reads for MAGNT S.14353-006. Reads were mapped to NCBI's non-redundant database with Diamond, and illustrated with MEGAN. Only nodes with a minimum support of 50 reads are shown.

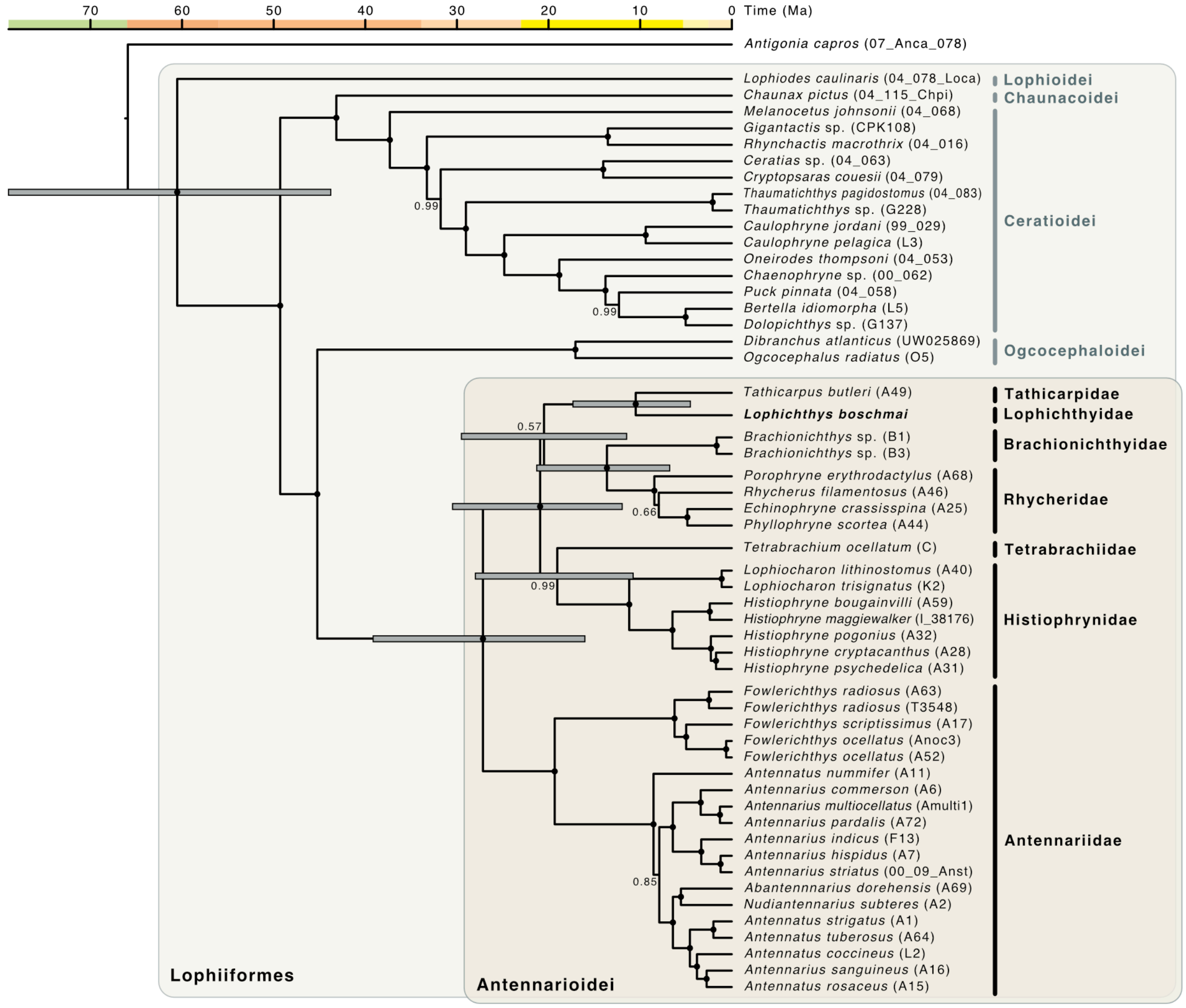


Supplementary Figure S4: Time-calibrated phylogeny reconstructed from genome-wide UCE marker sequences, after strict filtering for missing data (allowing a maximum of one missing nucleotide per site). The phylogeny was reconstructed by Bayesian inference with BEAST2. Black circles on nodes indicate full (1.0) Bayesian posterior probability; labels mark nodes with Bayesian posterior probabilities below 1.0. For the calibrated node (the first divergence among Lophiiformes) and for inter-familial divergences among Antennarioidei, node bars are shown to indicate 95% highest posterior density intervals for age estimates. The order Lophiiformes, lophiiform suborders, and families within the suborder Antennarioidei are highlighted.

SUPPLEMENTARY TABLES

| **Species** | **Individual ID** |
|---|---|
| *Antennarius commerson* | A6 |
| *Antennarius hispidus* | A7 |
| *Antennarius indicus* | F13 |
| *Antennarius multiocellatus* | Amulti1 |
| *Fowlerichthys ocellatus* | Anoc3 |
| *Antennarius pardalis* | A72 |
| *Fowlerichthys radiosus* | A63 |
| *Antennarius sanguineus* | A16 |
| *Antennarius striatus* | 00 09 Anst |
| *Antennatus coccineus* | L2 |
| *Abantennnarius dorehensis* | A69 |
| *Antennatus nummifer* | A11 |
| *Antennatus rosaceus* | A15 |
| *Antennatus strigatus* | A1 |
| *Antennatus tuberosus* | A64 |
| *Antigonia capros* | 07 Anca 078 |
| *Ceratias* sp | 04 063 |
| *Brachionichthys* sp | B1 |
| *Brachionichthys* sp | B3 |
| *Caulophryne jordani* | 99 029 |
| *Caulophryne pelagica* | L3 |
| *Bertella idiomorpha* | L5 |
| *Cryptopsaras couesii* | 04 079 |
| *Chaunax pictus* | 04 115 Chpi |
| *Chaenophryne* sp | 00 062 |
| *Dibranchus atlanticus* | UW025869 |
| *Dolopichthys* sp | G137 |
| *Echinophryne crassisspina* | A25 |
| *Fowlerichthys ocellatus* | A52 |

Supplementary Table S1: Individuals used in nuclear phylogenetic analyses of the UCE data of Hart et al. (2022).

| **Species** | **Individual ID** |
| --- | --- |
| *Fowlerichthys radiosus* | T3548 |
| *Fowlerichthys scriptissimus* | A17 |
| *Gigantactis* sp | CPK108 |
| *Histiophryne bougainvilli* | A59 |
| *Histiophryne cryptacanthus* | A28 |
| *Histiophryne maggiewalker* | I 38176 |
| *Histiophryne pogonius* | A32 |
| *Histiophryne psychedelica* | A31 |
| *Lophiocharon lithinostomus* | A40 |
| *Lophiocharon trisignatus* | K2 |
| *Lophiodes caulinaris* | 04 078 Loca |
| *Melanocetus johnsonii* | 04 068 |
| *Nudiantennarius subteres* | A2 |
| *Ogcocephalus radiatus* | O5 |
| *Rhynchactis macrothrix* | 04 016 |
| *Phyllophryne scortea* | A44 |
| *Porophryne erythrodactylus* | A68 |
| *Puck pinnata* | 04 058 |
| *Rhycherus filamentosus* | A46 |
| *Oneirodes thompsoni* | 04 053 |
| *Tathicarpus butleri* | A49 |
| *Tetrabrachium ocellatum* | C |
| *Thaumatichthys pagidostomus* | 04 083 |
| *Thaumatichthys* sp | G228 |

Supplementary Table S1 (continued): Individuals used in nuclear phylogenetic analyses of the UCE data of Hart et al. (2022).

| Position | Feature | Change | Comment |
|---|---|---|---|
| 383 | 12S | A → G | |
| 880 | 12S | T → C | |
| 2,361 | 16S | C → G | Probable error at end of assembled fragment, not inferred from reconstruction by mapping |
| 4,681 | ND2 | T → C | Synonymous |
| 11,199 | ND4 | C → T | Synonymous |
| 13,907 | ND6 | C → T | Nonsynonymous, Val replaced by Met |
| 15,638–15,674 | D-loop | 37-bp indel | |
| 15,699 | D-loop | A → G | Probable error at end of assembled fragment, not inferred from reconstruction by mapping |
| 15,705 | D-loop | T → C | |
| 15,725 | D-loop | 1-bp indel | Probable error at end of assembled fragment, not inferred from reconstruction by mapping |
| 15,743 | D-loop | T → C | |
| 16,134 | D-loop | C → T | |

Supplementary Table S2: Observed differences between the mitochondrial genomes of the two *Lophichthys boschmai* specimens. Changes are reported relative to S. 14545-001.

| Species | Sample | Museum specimen | Library name | Genbank accession (mito genome) | ENA study accession | ENA sample accession | ENA experiment accession | ENA run accession | Short read data reference |
|---|---|---|---|---|---|---|---|---|---|
| *Lophichthys boschmai* | NTM S.14545-001 | NTM S.14545-001 | lib-0185 | PP328847 | PRJEB83588 | ERS22579803 | ERX13467678 | ERR14064556 | this study |
| *Lophichthys boschmai* | NTM S.14353-006 | NTM S.14353-006 | lib-0192 | PP328846 | PRJEB83588 | ERS22579804 | ERX13467679 | ERR14064557 | this study |
| *Tathicarpus butleri* | A47 | WAM 32903.001 | NA | NA | PRJNA810755 | SAMN26280027 | SRX14368933 | SRR18222505 | Hart *et al.* (2022) |
| *Tetrabrachium ocellatum* | TR1 | UW049710_TR1 | NA | NA | PRJNA578585 | SAMN13066224 | SRX7031343 | SRR10320468 | Swann *et al.* (2020) |

Supplementary Table S3: Information on short-read sequence data
